# Myopic control of neural dynamics

**DOI:** 10.1101/241299

**Authors:** David Hocker, Il Memming Park

**Affiliations:** Department of Neurobiology and Behavior, Stony Brook University, Stony Brook, NY 11794; Department of Applied Mathematics and Statistics, Stony Brook University, Stony Brook, NY 11794; Institute for Advanced Computational Science, Stony Brook University, Stony Brook, NY 11794

## Abstract

Manipulating the dynamics of neural systems through targeted stimulation is a frontier of research and clinical neuroscience; however, the control schemes considered for neural systems are mismatched for the unique needs of manipulating neural dynamics. An appropriate control method should respect the variability in neural systems, incorporating moment to moment “input” to the neural dynamics and behaving based on the current neural state, irrespective of the past trajectory. We propose such a controller under a nonlinear state-space feedback framework that steers one dynamical system to function as through it were another dynamical system entirely. This “myopic” controller is formulated through a novel variant of a model reference control cost that manipulates dynamics in a short-sighted manner that only sets a target trajectory of a single time step into the future (hence its myopic nature), which omits the need to pre-calculate a rigid and computationally costly neural feedback control solution. To demonstrate the breadth of this control’s utility, two examples with distinctly different applications in neuroscience are studied. First, we show the myopic control’s utility to probe the causal link between dynamics and behavior for cognitive processes by transforming a winner-take-all decision-making system to operate as a robust neural integrator of evidence. Second, an unhealthy motor-like system containing an unwanted beta-oscillation spiral attractor is controlled to function as a healthy motor system, a relevant clinical example for neurological disorders.

## 1 Introduction

Advances in recording technology are making it possible to gain real-time access to neural dynamics at different length and time scales [BRAIN, 2014, Jun et al., 2017], allowing us to consider the structure of the brain’s operation in ways that were previously inaccessible. Central to that understanding of neural dynamics is the widely-held belief that *dynamical systems* underlie all of the core operations of neural systems [Breakspear, 2017, Fairhall and Machens, 2017, Sussillo, 2014, Izhikevich, 2006]. Dynamical systems are systems of time-independent dynamics that drive the evolution of a set latent states that may or may not be direclty observable, which in neural systems are proposed to account for motor function [Churchland et al., 2012], cognitive processes [Park et al., 2014, Mante et al., 2013, Jazayeri and Afraz, 2017], and sensory processing [Li et al., 2017]. The controlled stimulation of neural systems offers not only a novel tool to perturbatively study the underlying dynamical systems; but also shows tremendous potential to treat a host of brain disorders, ranging from movement diseases such as Parkinson’s disease and essential tremor [Deuschl et al., 2006, Lyons and Pahwa, 2004], epilepsy [Handforth et al., 1998, Morrell, 2011], and even mood disorders such as severe depression [Ineichen et al., 2016]. In particular, there has been recent success in combining real-time neural data acquisition with closed-loop stimulation for treating Parkinson’s disease [Rosin et al., 2011, Malekmohammadi et al., 2016].

Unfortunately, the current framework for manipulating neural systems is not structured to deal with the unique challenges posed by controlling complex neural dynamics. One of the central goals of control theory is to manipulate a system to mimic some or all characteristics of a target system of dynamics, and nearly all control systems accomplish this by controlling the system state to either track a specified target trajectory or to regulate to a known set point [Ioannou and Sun, 2012]. Closed-loop control systems specifically designed for neural systems also operate under this paradigm [Schiff, 2011, Yang and Shanechi, 2016, Newman et al., 2015], and clinical devices use even more simplistic open-loop or reactive protocols [Little et al., 2013, Rosin et al., 2011, Morrell, 2011]. If neural systems function as a dynamical system by nonlinearly filtering signals [Freeman, 1975, Haykin and Principe, 1998], then significant portions of the observed neural fluctuation would correspond to relevant exogenous input signals to the system such as volition, memory or sensory information. Such controls designed to move to or maintain a target state counteract any natural fluctuation in neural trajectories, and create a rigid system that is no longer dynamically computing. For example, when building neural prosthetics for an abnormal motor-related brain area, it is crucial for the controlled neural activity to be close to normal; however, simply controlling it to replay a fixed motor command would not allow flexibly changing one’s mind mid action. Therefore, any control objective that only considers externally set constraints through trajectory or set-point control would be limited both in their application for treating neurodynamic diseases as well as for studying neural computations in cases where preserving dynamic information processing capability is important.

Given this perspective, we propose a new control objective called *myopic control* that respects the unforeseeable variability in neural systems. The objective of myopic control is for the controlled system to behave as a target neural dynamical system. This is reminiscent of a well-developed field in control theory known as model reference control (MRC) [Ioannou and Sun, 2012], though MRC has been widely used for trajectory-tracking problems. Unlike MRC, myopic control is independent of the past trajectory and does not account for the far future—given the current state of the system, it tries to behave as the target dynamical system *instantaneously*.

The qualitative difference between our control scheme and trajectory-tracking methods is depicted in Fig. 1. Given some target dynamical system, utilizing trajectory control would force the neural system to follow a target trajectory, although not through the true target dynamics. Scenarios may arise where trajectory control and myopic control may be very similar (Fig. 1A), although there can be fundamental, qualitative differences in the presence of noise or large disturbances due to exogenous inputs (Fig. 1B). In that case, the trajectory resulting from trajectory control would not be generated from the target dynamics, and forces the state to evolve toward the pre-computed target state. In this way, our controller preserves the full neural variability of our target dynamics, ranging from potentially different trajectories towards the same fixed point to even allowing for potentially different behavior than expected.

**Figure 1:**
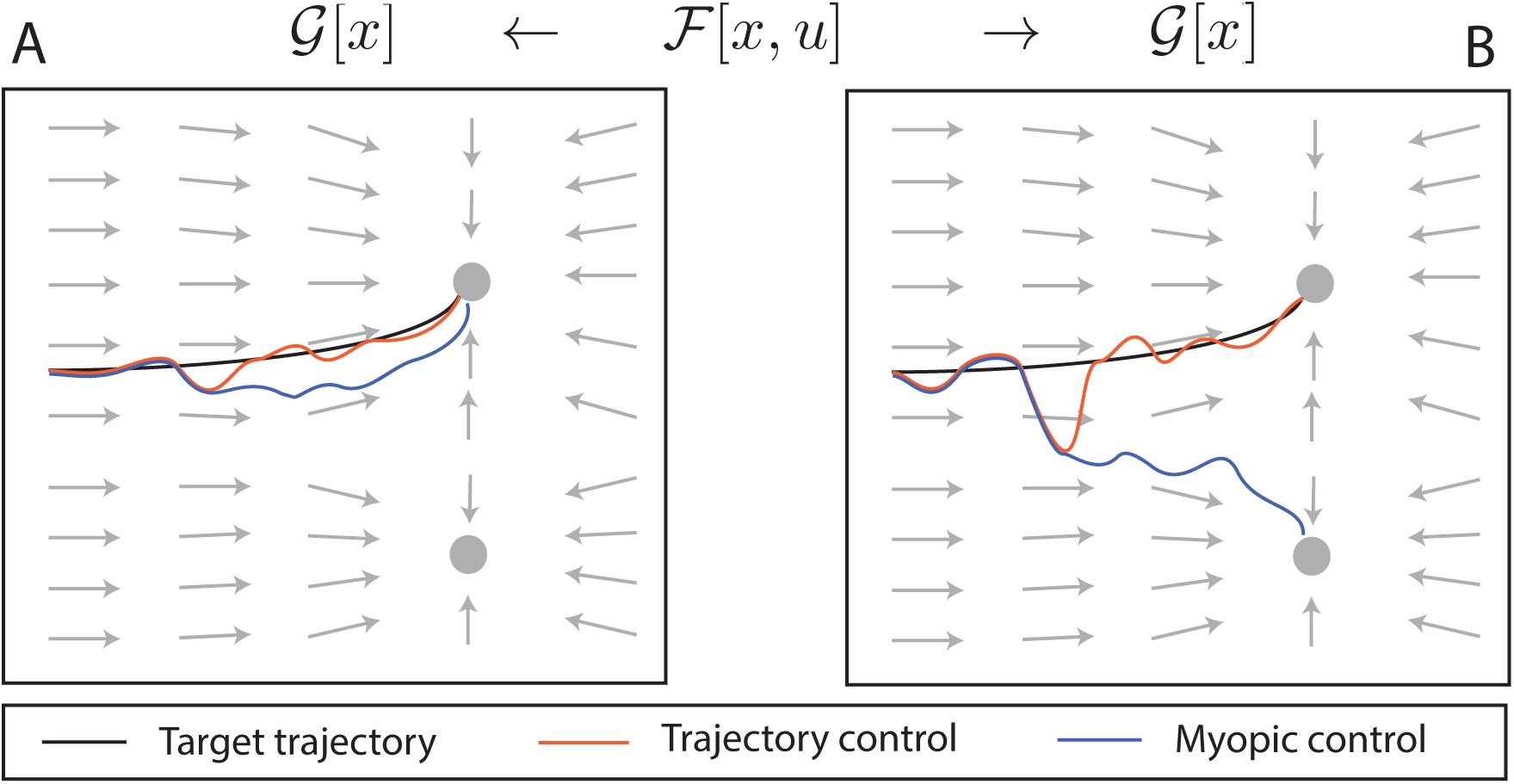
Qualitative difference between our proposed *myopic* control of dynamics and trajectory control. Here ℱ is controlled to perform an example target dynamics 𝒢 (*e.g.*, perform a motor command), where its gradient flow is given in gray and two attractors are denoted as circles. A precomputed target trajectory **x_t_** through 𝒢 is shown in black. **A**) In the presence of small disturbances, the evolution of trajectory control forces the system back to **x_t_**, whereas myopic control allows for natural deviations. **B**) A large disturbance away from **x_t_** corresponding to an exogenous input that changes the target attractor mid-trajectory could lead to entirely different behavior between the two control methods. Only myopic control would capture the response of this disturbance through the true dynamics of 𝒢, while trajectory control blindly follows **x_t_**.

The paper is organized as follows. First, we formulate the goals of our control objective for manipulating neural systems, then define myopic control for linear and nonlinear dynamics. Next, we discuss some design features of how to construct the target dynamics of a desired dynamical system, and what types of difficulties may arise when trying to define healthy or desired neural dynamics. We then demonstrate this control’s ability to make dynamical systems act as though they were another system entirely through two relevant examples. First, a winner-take-all decision-making model is transformed to operate as a robust neural integrator of information when shown a stimulus in a forced, two-choice decision-making task. Second, a “diseased” motor system containing an unwanted beta-oscillation state is controlled to function as a healthy motor system, which is a motivating example for the treatment of movement disorders or other diseases with an underlying neurological state.

## 2. Materials and methods

### 2.1 Myopic dynamics control

Here we discuss the control problem of utilizing a dynamical system to behave as a separate dynamical system. Using a Bayesian state-space modeling framework[Ogata, 2010], we are interested in the time evolution of a posterior distribution of time-dependent, *n*-dimensional (latent) brain state *x*_*t*_ that are governed by (stochastic) dynamics *ℱ*[*x*_*t*_, *u*_*t*_] *≡ℱ_t_* with an *m-*dimensional control signal *u*_*t*_,

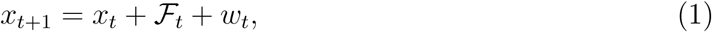

where *w*_*t*_ ∼ 𝒩(0, *Q*) is the state noise upon the dynamics. A second set of *target* stochastic dynamics 𝒢[*x*_*t*_] ≡ 𝒢*_t_* under which we would like our state to evolve, acts analogously on the state as

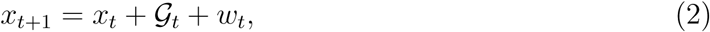

The noise in both dynamics is the same, as we are considering transforming ℱ into 𝒢 in the same physical neural system. In general, the control acts upon the dynamical system latent states *x* that may not be directly observable, and would need to be inferred from a set of observable variable to which the latent states are linked through an observation model. The influence of the controls would also be manifested in the observed states, though without loss of generality we have chosen to simplify our dynamics by omitting an observation model. These dynamics are in general nonlinear, and we denote their Jacobians (linearization at the current state and stimulus) as

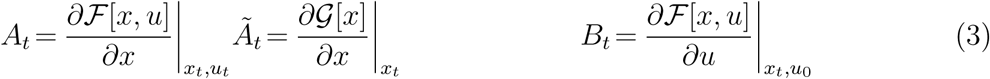

Arguably the most developed form of model-based control occurs for linear systems with quadratic costs on the state and control, known as linear quadratic gaussian (LQG) control [Stengel, 1994]. Finite-time horizon LQG controllers are optimal for costs of the simplified form

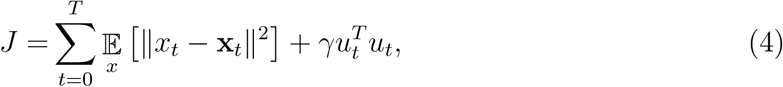

with linear dynamics ℱ_*t*_ = *Ax*_*t*_ + *Bu*_*t*_ + *w*_*t*_, and a regularization penalty factor *γ* is added onto the control power. The goal of minimizing (4) is to balance tracking along a target trajectory **x_t_** with the cost of implementing a control. The optimal LQG controller form 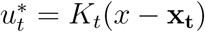 with gain *K*_*t*_ is found by solving the associated recursive Riccati equation from an end-point condition, and is a time-dependent controller through the time-dependence on *K*_*t*_ [Stengel, 1994].

Generating target dynamics is similar in spirit to LQG-type costs, although instead we are interested in minimizing the difference between the effect of target dynamics and controlled dynamics alongside control costs. Requesting that the controlled dynamics of *ℱ*_*t*_ act as through they are in fact *𝒢*_*t*_ can be written in a regularized, stepwise quadratic form as

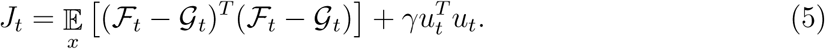

Note that this cost is defined at each time point *t*, and depends on the current state (posterior) distribution over *x*_*t*_. Utilizing control to track a defined trajectory that is generated from an uncontrolled set of target dynamics 𝒢 is the essence of model reference control (MRC), [Ioannou and Sun, 2012] although the costs associated with this control design are traditionally limited to regulation of a controlled trajectory around a set point or tracking of a predefined target trajectory evolving under 𝒢. Our cost in (5) instead seeks a control that effectively recreates a *single* step of a target trajectory from 𝒢, which to our knowledge is a major departure the typical use of model reference control. By weighting the difference between dynamics over a single time step, this myopic (*i.e.*, one-step) form negates the need to solve the Riccati equations, and the derivative *∂J/∂u_t_* can be straightforwardly calculated to identify the optimal *myopic control*.

Our work in this paper focuses primarily on designing a controller that optimizes eq. (5), which would be optimal for generating target dynamics over a single step. Since the controller would no longer contain any time dependence (the dynamics ℱ*_t_* and 𝒢*_t_* are indexed by their current time, but are dependent upon the state *x*_*t*_ only), it would generate a dynamical system with the same state space. Our controller does not assume the role of performing the bulk action on the state, which is instead encompassed in the original dynamics ℱ that presumably perform some form of related dynamics well. This is especially important in the context of neural dynamics performing a computation, where it would be undesirable for our controller to first perform the computation itself by tracing out a predefined trajectory **x_t_**. Instead, myopic control will assist that system’s natural ability to perform a neural computation.

The qualitative advantages of myopic control are depicted in Figure 1, in which the evolution of a trajectory-controlled system tracking a defined trajectory **x_t_** in a target dynamical system is compared to the evolution of a myopically controlled system designed to perform the target dynamics. In a noiseless environment, both trajectories would be identical; however, in the presence of small disturbances away from **x_t_**, tracking control would correct the trajectory in a distinctly non-dynamical fashion, evolving not through 𝒢 but instead forcing the system back onto **x_t_** in an unnatural manner (Fig. 1A). Myopic control would instead lead trajectory through the natural dynamics of 𝒢, which may lead to the same stable point, but through a distinctly different trajectory. Some disturbances may lead to different behavior between the two control methods, though. Figure 1B shows this scenario, in which a disturbance is corrected by trajectory control back toward **x_t_**, while myopic control followed the flow of, which lead it to a different attractor point. If this target dynamics were a decision-making computation, for example, myopic control may have lead to a “wrong” decision; however, allowing a controlled neural system to operate imperfectly in the perspective of modern control is precisely the type of flexibility that should be achieved to maintain its natural operation.

In the following sections we derive the form of our myopic controller. Ideally, the controller formulation will be distinct from the state estimator providing the feedback signal, and leads us to consider variants of the controller that rely upon different moments of the underlying state distribution. We first begin with the case of linear dynamics to demonstrate the simplified form of myopic control and its properties, then move the more applicable nonlinear case.

#### 2.1.1 Linear dynamics

Here we demonstrate that the myopic controller for linear dynamics depends only upon the mean of the state distribution, and thus the state estimator and controller design are separable for myopic control.

**Theorem 1.** If target and controlled dynamics are linear in state *x* and control *u*, then myopic control depends only upon state mean 𝔼[*x*].

*Proof.* Let the linear dynamics under control and the target dynamics be

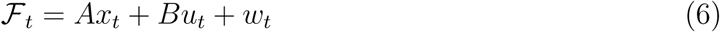

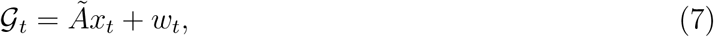

where the state distribution over *x* has first and second central moments 𝔼[*x*_*t*_] = *µ*_*t*_, 𝔼[(*x*_*t*_ − *µ*_*t*_)^2^] = ∑*_t_*, and the state noise is normal with *w*_*t*_ ∼ *𝒩*(0, *Q*). Expanding the dynamics cost in (5) gives

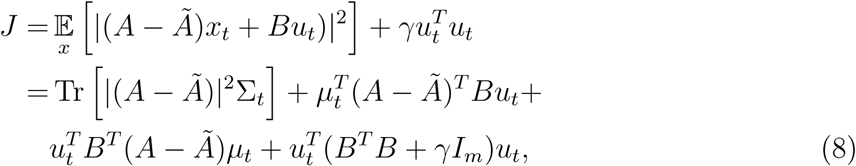

where *I*_*m*_ is the *m* × *m* identity matrix. By examining (8) it is clear that regardless of the distribution over *x*, the cost depends only upon the first two moments of the distribution of *x*. Maximizing (8) yields the optimal linear myopic controller form 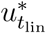, which depends only upon the state mean,

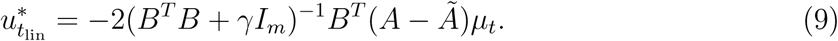

#### 2.1.2 Nonlinear dynamics controller with a moment expansion approximation

For nonlinear dynamics, simply differentiating (5) leads to an ill-suited expression for a controller, since there is an implicit dependence of the controller upon itself through ℱ. One approximation to alleviate this is to expand the nonlinear dynamics about null control (*u*_0_ = 0) to first order, with the form

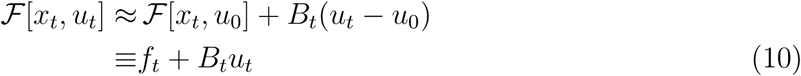

where *f*_*t*_ ≡ *f* [*x*_*t*_] ≡ ℱ[*x*_*t*_, *u*_0_] and for the remainder of the work *B*_*t*_ ≡ *B*[*x*_*t*_, *x*_0_] is the Jacobian of ℱ as in eq. (3). This leads to an expression for the derivative of *J* and myopic controller as

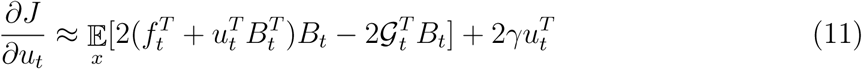

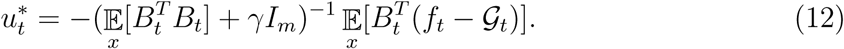

The expectations in (12) depend upon the state distribution of *x*_*t*_, although it would be desirable if akin to LQG that the controller was separated from state estimation, and only depended upon low-order moments of *x*. To construct such a controller we will expand 𝔼*_x_*[·] in terms of the mean and covariance of *x*_*t*_; in general, the terms in this expansion will contain Jacobian matrices, higher order derivatives, and state vectors that are all evaluated at the distribution mean *µ*_*t*_, multiplied by the covariance ∑*_t_* in some form. For example, the Jacobian *B*_*t*_ is expanded as

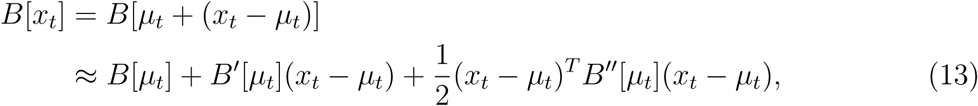

and would follow similarly for the other terms in 𝔼_*x*_[·]. Such an approximation is valid when the deviations from our estimated state *µ*_*t*_ are small, and in this regime only low-order moments are necessary. It is assumed that state estimation to obtain *µ*_*t*_ and ∑*_t_* can be performed regularly enough in practice to operate in the regime such that (13) is valid, and we will consequently consider two forms of nonlinear myopic control. *First-order* myopic control will include only terms dependent upon state mean, just as in the linear dynamics case of the previous section. *Second-order* myopic control will analogously depend upon both *µ*_*t*_ and ∑*_t_*. In each controller the terms *f, 𝒢, B* and derivatives 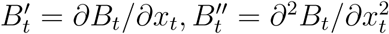, will all be evaluated at the distribution mean *µ*_*t*_ and null control *u*_0_ = 0, so we will temporarily drop the functional dependence of these terms in the notation. The prime notation will indicate a derivative with respect to state.

The first expectation in (12) includes only terms relating to *B.* Expanding and keeping terms up to second order gives

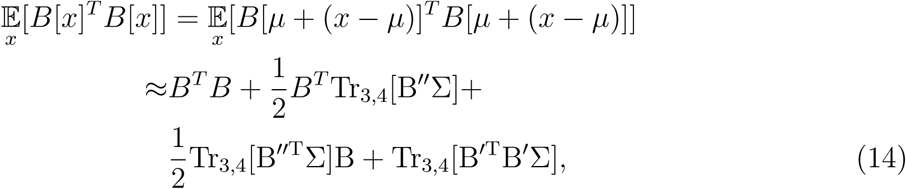

where Tr_3,4_ denotes the partial trace over dimensions 3 and 4. For an (*n × m × n × n*) tensor *T* this operation maps to an (*n × m*) matrix *M* = Tr_3,4_[T] as

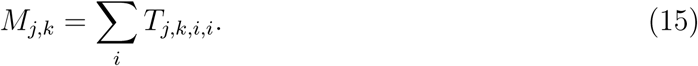

Similarly, expanding 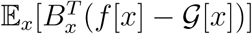 up to second order yields

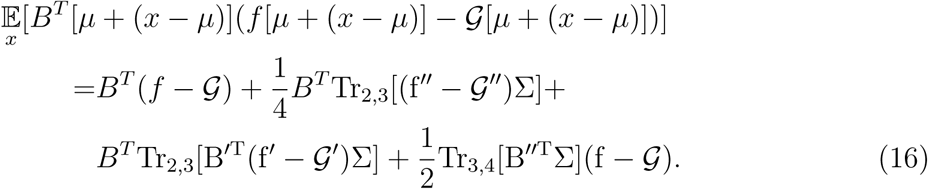

(12), (14), and (16) define our second-order nonlinear myopic controller *u*_2nd_, and simply omitting the covariance-dependent terms gives first order controller expansions,

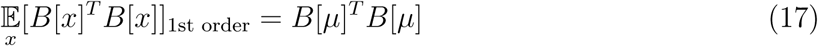

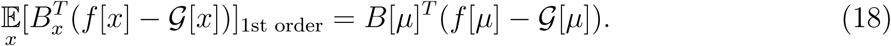

First-order control *u*_1st_ is attractive for its simplicity, and it is important to ask under what circumstances would first-order control outperform second-order control? First, if Tr(∑*_t_*) is very small (*i.e.*, small uncertainty about the state *x*_*t*_), then second-order terms are negligible. Second, by noting that nearly all second-order terms contain derivatives of *B, f* and, another regime in which first-order control may be superior is under “super smooth” dynamics in which the magnitude of successive derivatives is smaller than the previous one (*e.g. ||B|| > ||B′|| > ||B″||*). Moreover, if the control portion of the Jacobian *B* is state-independent, then second-order control only has one covariance-dependent term in 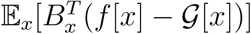,

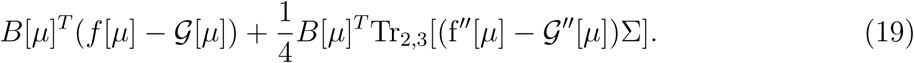

### 2.2 Evaluating controller performance

The performance of a myopic controller is formally benchmarked by the regularized cost in (5), although it is important to isolate a cost describing the performance of only the dynamics. The cost of expected mean performance is denoted by 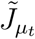, and is given by

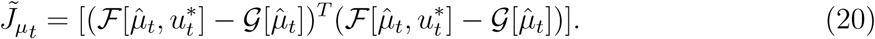

This cost is easy to compute and provides information about the mean behavior of the control, although it ignores the variability of the state distribution ∑*_t_*. A more informative cost incorporates the impact of the entire distribution of *x*, denoted by 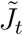 as

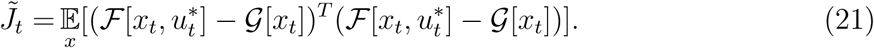

We estimate (21) through Monte Carlo integration assuming the maximum entropy distribution at each time point given the first two moments, *i.e.*, a normal distribution 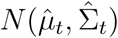.

### 2.3 Design principles for targeted dynamical systems

Myopic control omits the requirement of supplying a target neural trajectory or set point in the neural state space, which resonates with our design requirement of a future-agnostic controller that need not prescribe what the brain should be doing precisely.

Balancing the simplicity and ease of myopic control, though, is the relative complexity in designing a target dynamical system 𝒢*_t_*. At first glance, it may seem as though we have merely shifted complications of controlling neural dynamics. However, this perspective more clearly frames the goal of neural dynamics control, and we believe that it identifies a general design question yet to be seriously considered by the neural processing community: Given a rough sketch of neural dynamics and a desire to change them, what is an appropriate target dynamical system?

The choice of 𝒢 can roughly be broken down into three design problems for dynamical systems: i) removal or avoidance of an unwanted feature, ii) addition of a desired feature, and iii) modification of an existing feature. For example, there may be attractors (representing macrostates) in ℱ indicative of a dysfunctional behavior that should be avoided for healthy brain function, such as limit cycle attractors. Or, one may wish to introduce additional attractor macrostates in a decision-making system in order to support robust neural integration of evidence [Koulakov et al., 2002]. We will consider both of these scenarios in the following sections. Our ideal design approach used here is summarized in Figure 2, which is to use multiplicative filters upon the controlled dynamics ℱ to preserve desired features, with the addition of either barrier functions to remove undesirable aspects of ℱ or to prevent access into that region of state space. Alternatively an additive function could be utilized to introduce new features. Care must be taken with the shape and positioning of the additive barrier or extra feature though, as any zero crossings of this additive term will introduce fixed points into the dynamics. In the example case in Figure 2, a barrier function is used to remove an undesirable feature of the dynamical system by producing a net rightward gradient flow in the low *x*_1_ region of state space, where the zero crossing of the barrier function is aligned with other fixed points of the system that are denoted in blue. Under this strategy, we can also view modification of an existing feature as simply a removing it and replacing it with the desired one.

**Figure 2:**
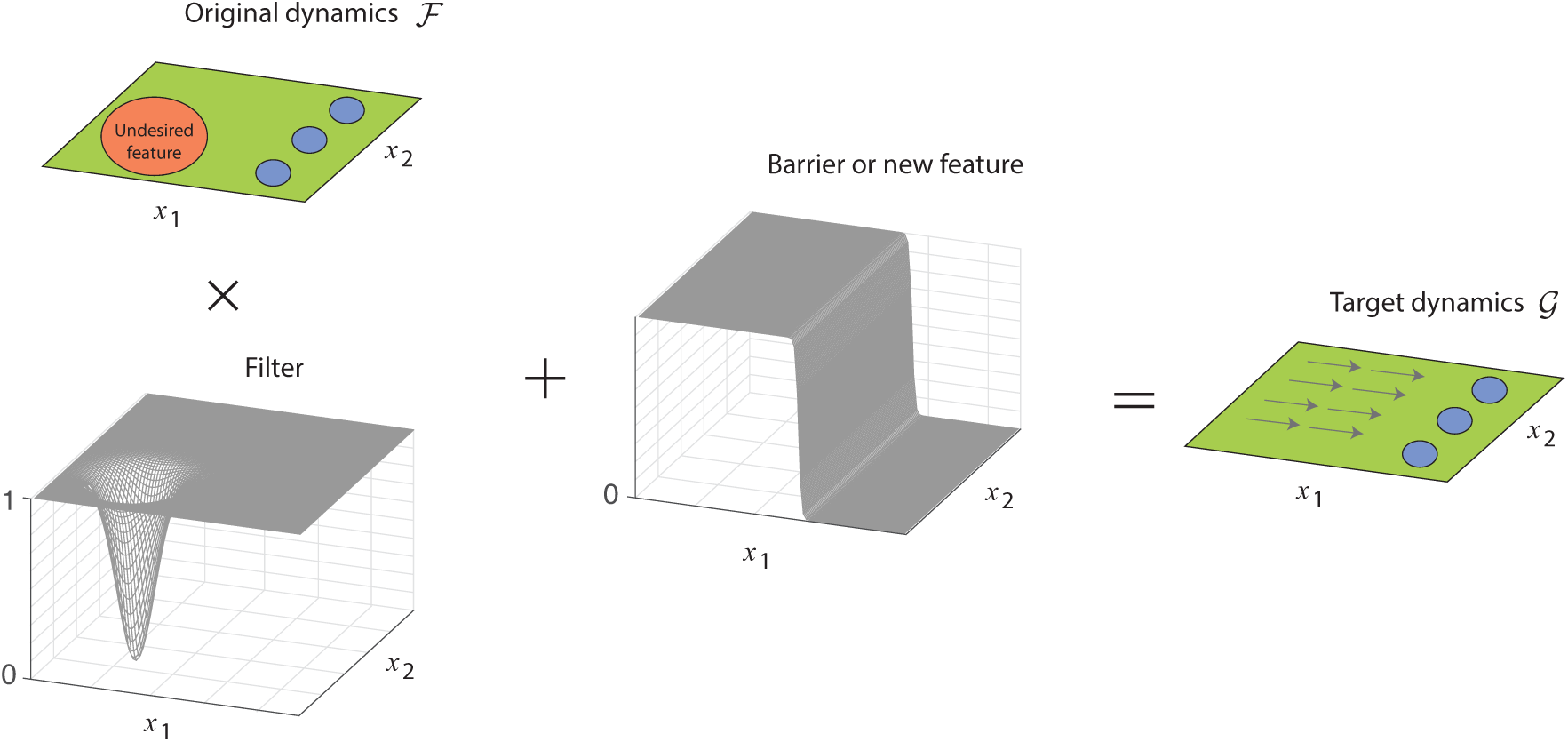
Design strategy for creating target dynamics 𝒢. Green and blue regions represent features of the original dynamics ℱ that are to be maintained, while an undesirable feature is denoted with orange. A multiplicative filter removes the unwanted feature, while an additive barrier function prevents access to unwanted state space by enforcing a gradient flow toward desirable regions with well-behaved dynamics.

### 2.4 Numerical methods

In the follow sections we detail the dynamics of each dynamical system in the examples. Since the primary objective of this work is to understand the performance of the myopic controller, in both examples we used a simple state estimator to calculate 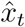 and 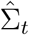, employing extended Kalman filtering (EKF) within Tensorflow assuming a noisy observation of the state as *y*_*t*_ = *x*_*t*_ + *v*_*t*_, where *v*_*t*_ ∼ *N* (0, *R*) and *R* is a diagonal covariance matrix. For lags in observations and control signal calculation, state prediction was performed by propagating 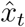 via,

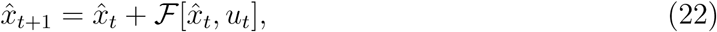

and covariance was estimated through sampling of time-evolved state predictions 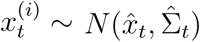 that also evolve in time using (22),

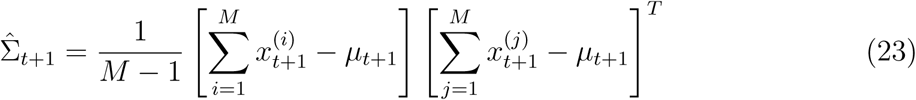

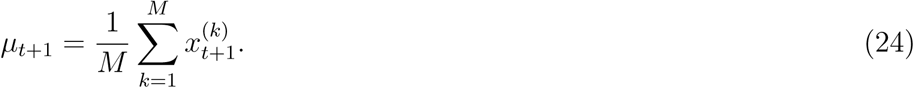

#### 2.4.1 Beta oscillation disease states

The diseased state dynamics are a modification of a dynamical system used to describe linear integrate-and-fire neurons as a limit cycle attractor [Izhikevich, 2006]. The specific limit cycle attractor is based upon the post-saddle-node bifurcation behavior at large current in the *I*_Na,p_ + *I*_K_ model from Ch. 4 of [Izhikevich, 2006] (eqs. 4.1-4.2). The high-threshold parameters in [Izhikevich, 2006] were utilized to generate the attractor, and the external current was tuned to generate a beta oscillation. The beta oscillation dynamics of state *X* = [*X*_1_, *X*_2_]*^T^* (omitting state and observation noise, for succinctness) are

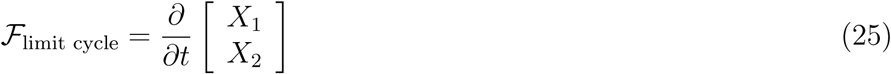

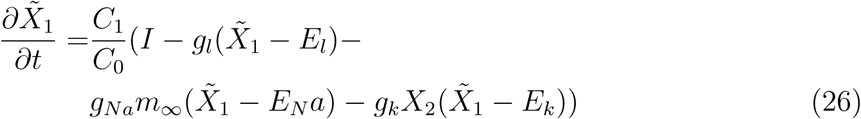

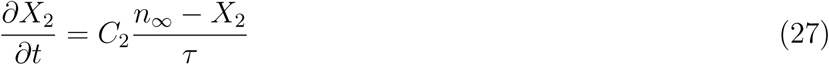

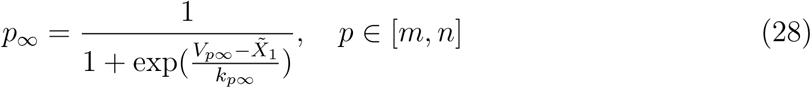

with parameters *C*_0_ = 1, *I* = 10, *E*_*l*_ = *-*80, *E*_*Na*_ = 60, *E*_*k*_ = *-*90, *g*_*Na*_ = 20, *g*_*k*_ = 10, *g*_*L*_ = 8, *τ* = 1, *V*_*m,∞*_ = − 20, *V*_*n,∞*_ = − 25, *k*_*m∞*_ = 15, *k*_*n∞*_ = 5. The original dynamics for *X*_2_ corresponded to an activation variable in an integrate-and-fire model, and as such were scaled to operate at the order of magnitude *X*_2_ *∈* [0, 1]; however, *X*_1_ was originally a voltage variable, and we rescaled it such that 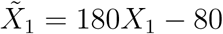. The magnitude of these dynamics were also scaled with *C*_1_ = 0.88 and *C*_2_ = 160 to reflect this change in *X*_1_.

The three stable points of the dynamics were added to the limit cycle attractor dynamics as two sets of gaussian-weighted Gabor functions centered at the three stable points *m*_1_ = [0.9, 0.25], *m*_2_ = [0.9, 0.50], *m*_3_ = [0.9, 0.75] with a width of the gaussian envelope *L* = 0.2. This portion of the dynamics was structured and scaled as

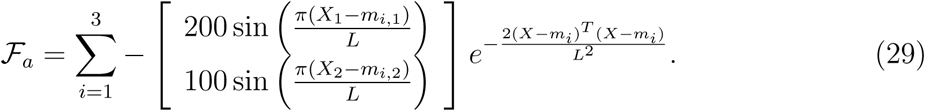

To combine the stable attractors and limit cycle attractors in a smooth fashion we adopted our design strategy of filtering out regions of the limit cycle attractor in the stable attractors regions around *m*_1_, *m*_2_, and *m*_3_, and added in the stable attractors. Finally, the state-independent control signal was added linearly to give the controlled dynamics with Δ*t* = 10^-4^*s* as

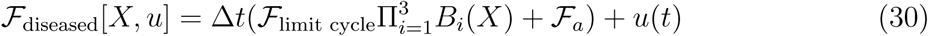

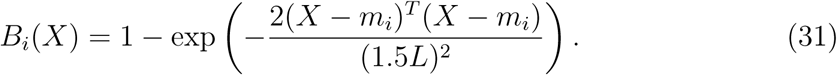

The healthy dynamics were designed by using the approach of Section 2.3 to encourage the dynamics to stay near stable attractors, and avoid the limit-cycle attractor. We designed a hyperbolic tangent filter function *FF* preserve the stable points, and a barrier function ℬ to encourage state movement away from the limit cycle

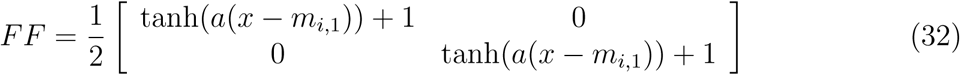

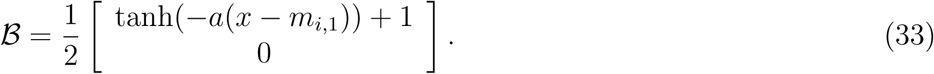

Both the filtering function and barrier function have their zeros at the intersection of the stable points, to avoid introducing additional unwanted stable points. The scale factor *a* = 20*π* creates a steep barrier. The healthy dynamics are then calculated by filtering on null-controlled unhealthy dynamics as

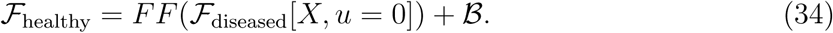

The state noise covariance *Q* = 10^*-*5^*I*_2_ was chosen to allow for noise-assisted departure of the uncontrolled dynamics from the stable points an into the limit cycle attractor. Observation noise covariances *R* ∈ [10^*-*6^*I*_2_, 10^*-*5^*I*_2_, 10^*-*4^*I*_2_] were used.

#### 2.4.2 Winner-take-all and robust neural integrator dynamics

The Winner-take-all dynamics are based upon the state-space description in [Wong and Wang, 2006], in which two sub-populations of excitatory neurons *X*_1_ and *X*_2_ have a reduced-state dynamical description for decision-making of the direction of a random moving dot visual stimulus. The dynamics for the two-dimensional state driven by control signals *u*(*t*) = [*u*_1_(*t*), *u*_2_(*t*)]*^T^* are given by

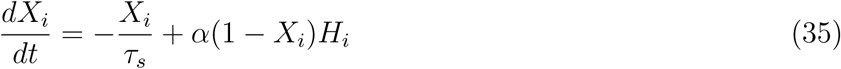

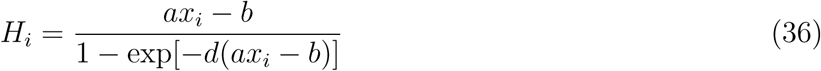

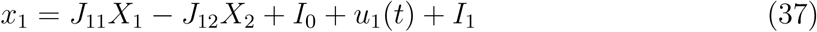

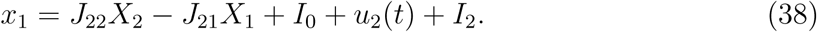

The visual stimulus is represented as input current *I*_1_ and *I*_2_ to each population with stimulus strength *µ*_0_ = 30*Hz* and directional percent of coherence *c′*,

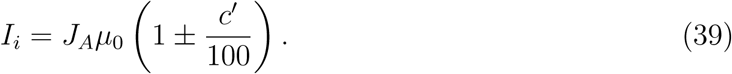

High activity of a state *X*_1_ corresponds to decision due to activity of that sub-population of neurons with positive-signed coherence of the stimulus, and *X*_2_ alternative has high activity for negative-sign coherence of the stimulus, indicating the direction of the stimulus. Parameter values reported in [Wong and Wang, 2006] were used. The controls to the system *u*_1_(*t*) and *u*_2_(*t*) were modeled as additional input currents to the sub-populations.

The target neural dynamics of a robust neural integrator are based conceptually upon [Koulakov et al., 2002], and their state space description is modeled as a set of hyperbolic tangents that generate interwoven nullclines. The two states *X*_1_ and *X*_2_ have gradients given by

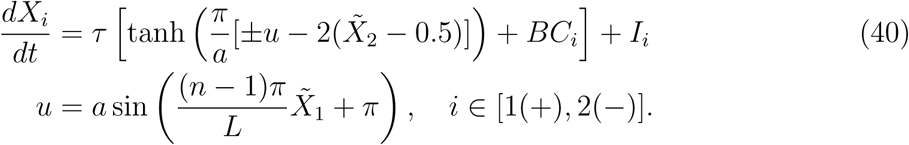

The nullcline shapes *u* are defined by *n –* 1 nodes over a length *L*, and have a hyperbolic tangent on each side in state space. (40) use a rotated set of state-space coordinates 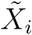 given by a rotation matrix 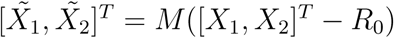, where *M* rotates by the angle *π/*4 and *R*_0_ = [*L/*2, 1]*^T^.* The boundary conditions *BC*_1_ and *BC*_2_ enforce the final fixed points of the neural integrator line to be global attractors, and are given by additional hyperbolic tangents of the form

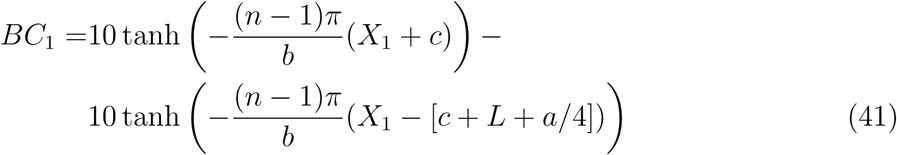

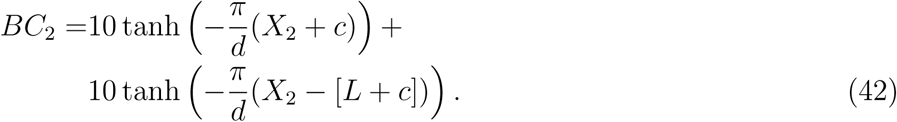

The parameters of the model were chosen to roughly match the magnitude and fixed-point locations of the winner-take-all dynamics. *τ* = 1*e*^-3^ (*i.e.*, 1ms timesteps), *a* = 0.2, *n* = 7, *L* = 0.7, *b* = 4*/*3, *c* = 0.083, *d* = 1.2. The stimuli to the robust neural integrator *I*_1_ and *I*_2_were given by

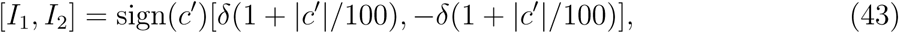

where δ = 7.5*e –* 4. The state noise covariance *Q* = 5 × 10^*-*5^*I*_2_ was chosen to allow the robust neural integrator to utilize the state noise to transition from one stable point to another, and observation noise covariances *R* ∈ [10^*-*6^*I*_2_, 10^*-*5^*I*_2_, 10^*-*4^*I*_2_] were used.

## 3 Results

In the following examples we demonstrate the ability of myopic control to match the dynamics of several relevant dynamical systems for neural computations. Simulations to benchmark the performance of myopic control were conducted using Tensorflow (Python API). System details can be found in the methods section, and code for the myopic controller is available at https://github.com/catniplab/myopiccontrol.

### 3.1 Robust neural integration from winner-take-all dynamics

We first deal with controlling neural computations for decision making, and demonstrate how myopic control can be used to change a winner-take-all (WTA) decision-making dynamics and convert it into a robust neural integrator (RNI). WTA dynamics for a simple, forced two-choice decision-making process function through a dynamical system where stimulus modulates the dynamics to flow toward one of two stable attractors. As time progresses, the neural state is driven toward one of the two stable attractors, each comprising a separate decision. In contrast, a robust neural integrator has multiple fixed points in between those two final stable attractors that allow for a stable, intermediate representation of accumulated evidence— creating robustness against uncertainty in stimulus and small internal perturbations.

We implemented a well-known approximation of a WTA dynamical system underlying two populations of spiking, excitatory neurons connected through strong recurrent inhibitory neurons, and our control for this system is an external injected current into each excitatory population [Wong and Wang, 2006]. Our target dynamics embody a low-dimensional analogue of the robust neural integration model suggested by Koulakov and coworkers [Koulakov et al., 2002]. Our RNI dynamical system is conceptually quite simple: Two sinusoidal nullclines that are interwoven can generate alternating stable and unstable fixed points, and with the addition of boundary conditions on the final stable fixed points can generate a dynamical system with a line of stable fixed points. The phase portraits for each system are shown in Figure 3.

**Figure 3:**
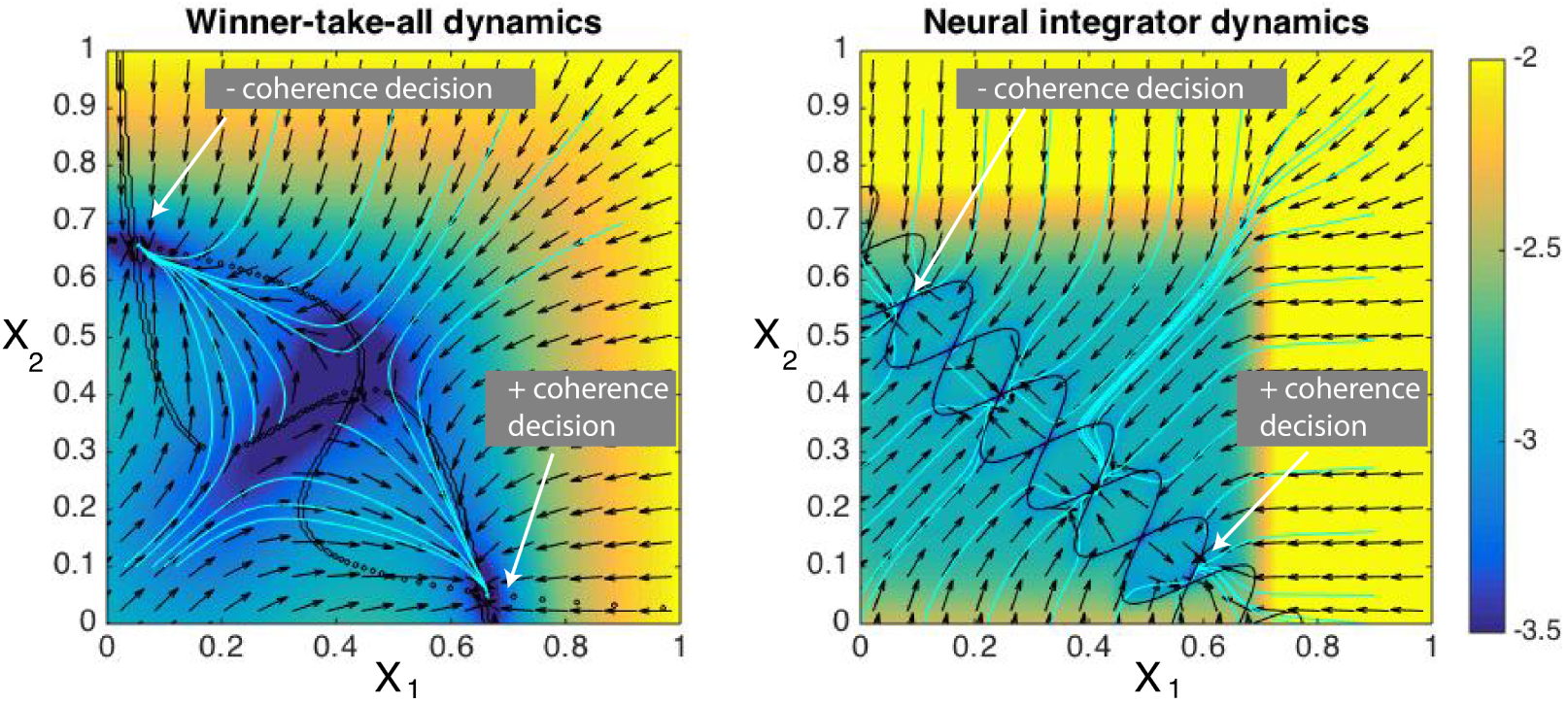
Phase portraits for the A) winner-take-all dynamics and B) robust neural integrator. The magnitude is plotted on a logarithmic scale for easier visualization, and arrows give gradient direction. Streamlines are depicted in cyan. The nullclines of each dynamical system are shown in black. The RNI system is formulated as an extension of the tangling of nullclines in the WTA dynamics, where additional crossings of the nullclines result in stable points.

A comparison between first-order myopic control and a trajectory control approach (*i.e.*, a control optimized for (4)) is presented in Figure 4. Specifically, we show that trajectory control possess shortcomings when dealing with particular decision-making tasks. Trajectory control approaches have an additional hurdle above myopic control in that there must be some policy in place to decide to which target **x***_t_* should evolve. The only way that trajectory control can help the neural system make an informed decision is by integrating the stimulus, itself, which assumes a role in the neural computation. Here, we allow the controller to observe and integrate 20ms of coherence at the beginning of the trial, and then use that information to prescribe a target point in state space (either the final + or – coherence decision points in Fig. 3). In this simulation the time-varying stimulus *c′* is initially uncertain, beginning with a small positive coherence for 500 ms and then changing to negative coherence for 500 ms, finally settling to a stronger negative coherence of *c′* = –12% for the remainder of a 1s trial, shown inset in Fig. 4. Both the target dynamics of robust neural integrator and myopic control to mimic it can adequately handle this “change-of-mind” in stimulus and eventually evolve its neural state to the negative coherence choice, but trajectory control system instead incorporates only the initial stimulus to incorrectly choose the positive coherence choice. Furthermore, it then holds it there with control, in spite of receiving new stimulus information that in some cases could have even been caught in the WTA system (Fig. 4, green).

**Figure 4:**
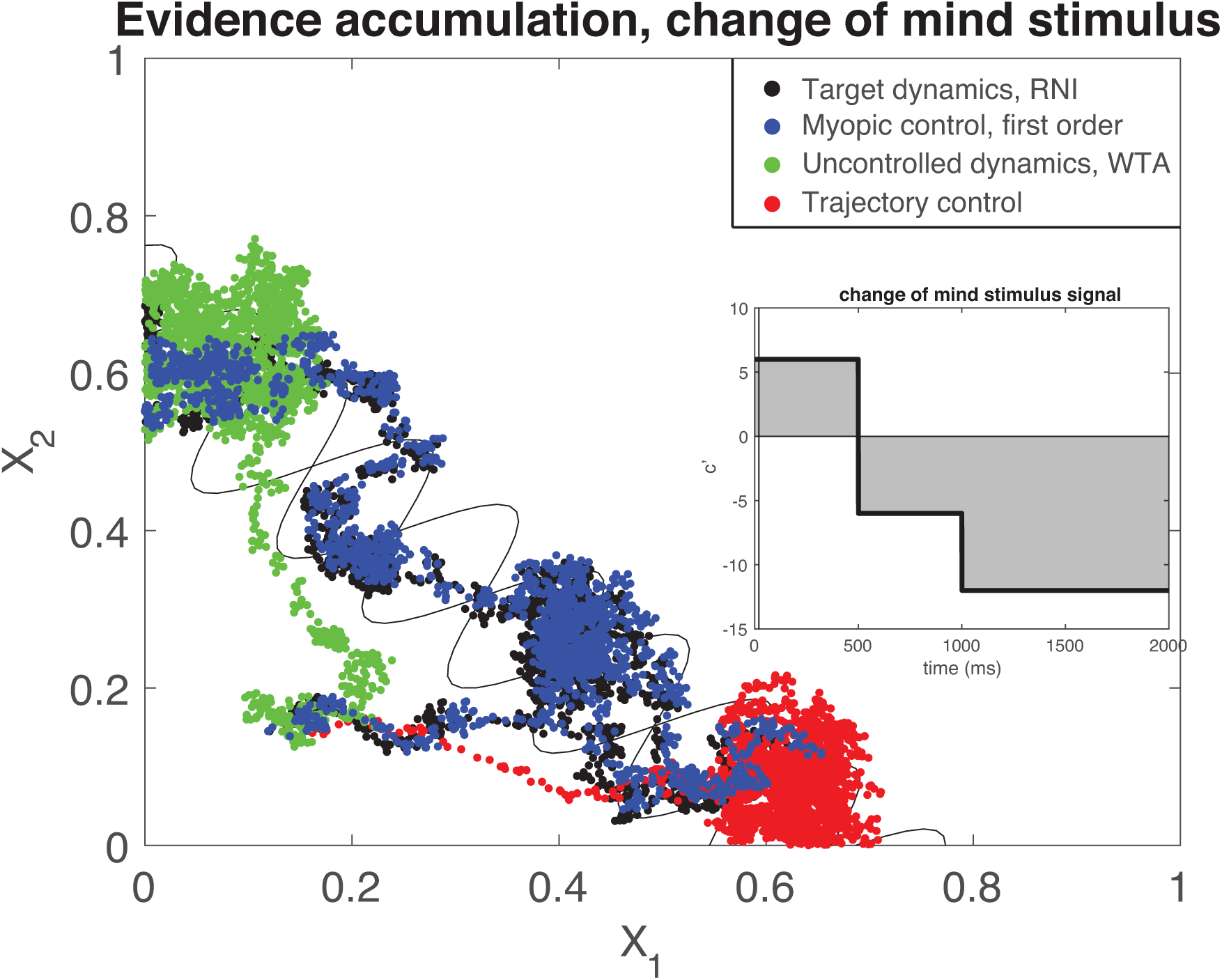
Example trajectories of controlled dynamics for an initially uncertain stimulus (inset). Trajectory controls with a policy to decide which decision to make by integrating the initial few ms of coherence signal not only steal the role of computational integration from the dynamical system, but in the presence of an initially uncertain stimulus perform poorly by making the wrong decision (red) where even a WTA system (green) can somewhat cope with a changing stimulus. Myopic control (blue) can integrate the incoming information just as the target RNI dynamics (black). inset: coherence for the change of mind stimulus. a final decision would be made by integrating the signal, signified by the gray region. Vertical line at 20ms indicates the portion of the signal integrated to decide on a target point for trajectory control.

An intuitive way to compare the performance of trajectory control vs. myopic control for decision making is to count the number of correct decisions made, which is summarized in Table 1, alongside the total power of the controls, calculated as

**Table 1:**
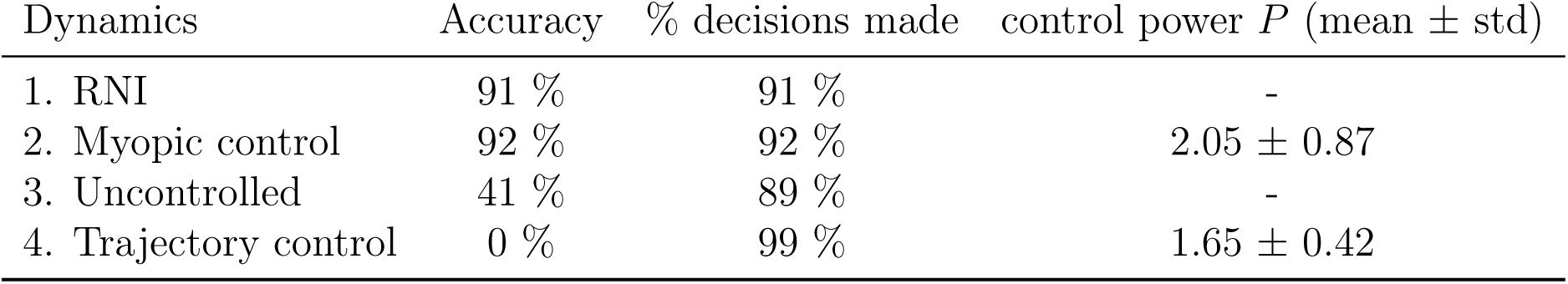
Accuracy of decision making for an uncertain stimulus.

| Dynamics | Accuracy | % decisions made | control power $P$ (mean ± std) |
| --- | --- | --- | --- |
| 1. RNI | 91 % | 91 % | - |
| 2. Myopic control | 92 % | 92 % | $2.05 \pm 0.87$ |
| 3. Uncontrolled | 41 % | 89 % | - |
| 4. Trajectory control | 0 % | 99 % | $1.65 \pm 0.42$ |

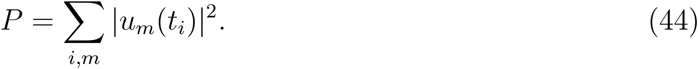

Accuracy for each control type calculated as percentage of correct fixed points chosen (noted in Fig. 3), and percentage decided as number of trials in which the state evolved to within a close radius of a decision point (radius = 0.15). Examining the accuracy of each method in the table, it is clear that this is an extreme case of how poorly trajectory control can perform, where even uncontrolled dynamics sometimes was able to change its mind and choose the correct stimulus. This is a specific example of our qualitative arguments against trajectory control that were shown in Fig. 1, in which markedly different behavior can be artificially enforced. Moreover, the total power required by the control signals is comparable, indicating that myopic control did not require substantially more power to perform the target dynamics

Quantitative performance of myopic control of a sample of 500 trials is summarized in Figure 5. Sample trajectories for uncontrolled, first-order controlled, and healthy dynamics are shown in Fig. 5A under the influence of an increasingly stronger time-dependent stimulus, denoted by coherence *c′*. Both the RNI and controlled system linger at an intermediate stable nodes before coherence has increased enough to make a more informed decision, indicated by the progression to the decision node. In contrast, the uncontrolled trajectory evolves straight to the decision without any intermediate stability at low coherence.

**Figure 5:**
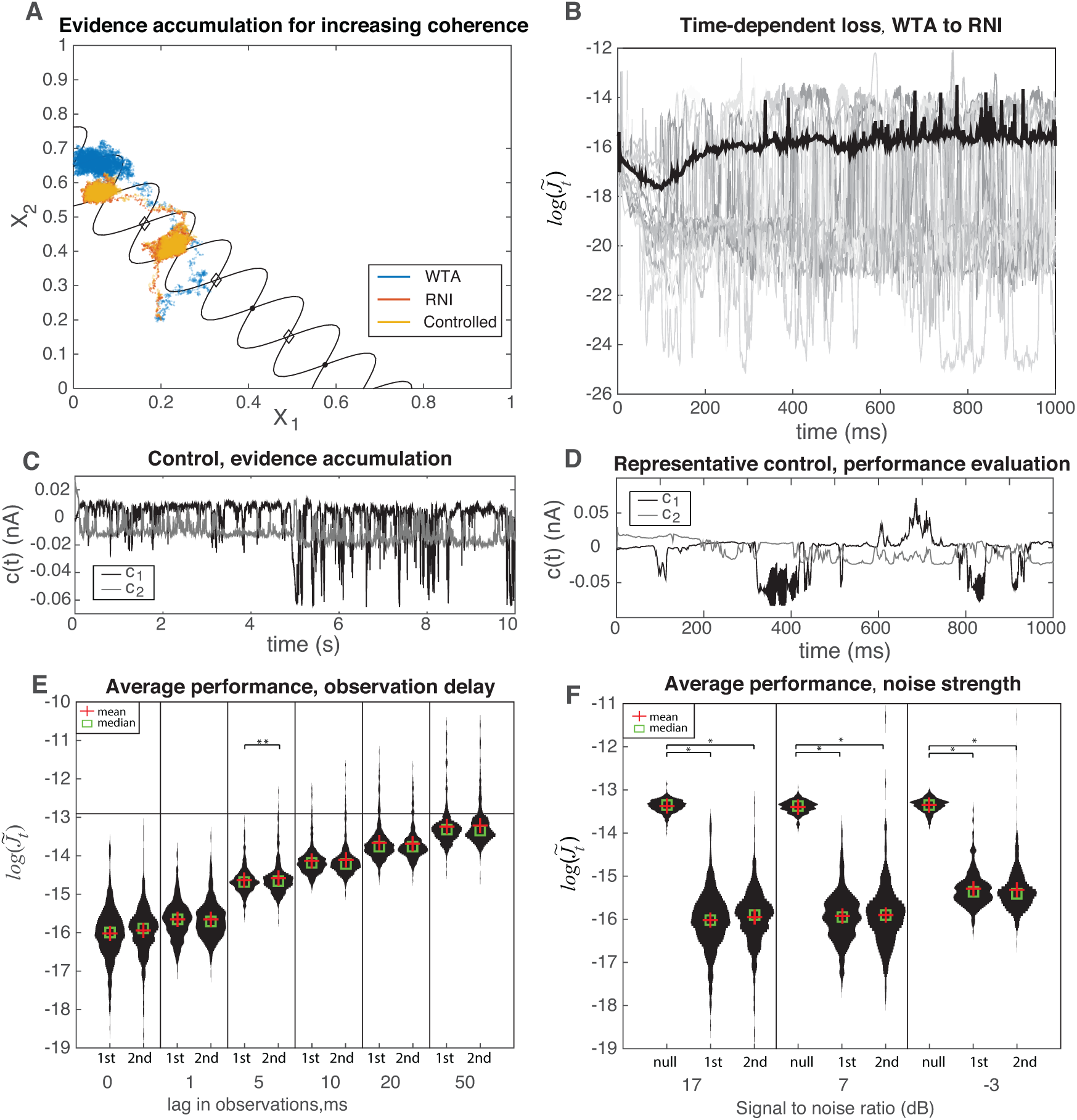
A) Example trajectory of controlled dynamics for evidence accumulation. Nullclines and stable points (dots) of the neural integrator shown in black, with unstable nodes shown as diamonds. B) Average time dependence of log-performance of expected cost in eq. (20) for winner-take-all to robust neural integrator dynamics with first-order control, SNR of 7dB, and fixed coherence *c′* = –6%. Example trials are shown in gray, and trial-averaged mean plotted in black. C) Control signals during the evidence accumulation with first-order control. D) Similar control signals for fixed-coherence trial. E) Violin plots showing distribution of time-averaged, log-performance for a lag in observations. F) Similar violin plots as in E), but for for varying observation noise strength, shown by signal-to-noise ratios (SNR) of system noise to observation noise, and including null control (*c*_*t*_ = 0). In both E) and F) * indicates *p <* 0.001 for a two-sample t-test. Unless otherwise noted, samples are not significantly different. **: *p* = 0.009.

Importantly, the controlled dynamics demonstrate the intermediate stability behavior found in robust neural integrators. Figure 5B summarizes the log-cost of 500 trials of first-order control with a fixed stimulus coherence of *c′* = –6%, where prototypical trials are shown in gray alongside the trial average in black. The control signals for the increasing coherence demonstration (Fig. 5A) and for the benchmark trajectories (Fig. 5B) are plotted in 5C and 5D, respectively. Again, promising and modest control amplitudes are observed in both cases. Finally, figures 5E and 5F show the time-averaged, log-performance for varying observation lag and noise strengths. Comparable to the previous section we see that second order control performs equivalently to first order across increasing observation lag and noise. However, the time-averaged distributions at low observation lag have quite long-tailed, unimodal distributions, and have negligible performance at a lag of 50 steps (note that Δ*t* is an order of magnitude higher for this system, which corresponds to a 50ms-ahead prediction). There is some change to a bimodal distribution for increasing observation noise in this system, but the notable feature is the increasingly long distribution tail for second-order control, which gives the opportunity for inferior performance as compared to first-order control.

### 3.2 Avoiding beta-oscillation disease states

Here, we aim to preserve an original set of dynamical features in ℱ while avoiding an unwanted regime of state space containing undesirable dynamics. This paradigm can act as the basis of state-space control for neurological disorders, where regions of state space may be associated with disease symptoms [Watter et al., 2015, Little and Brown, 2012]. Utilizing myopic control as a therapy for neurological disorders lends itself to considering which features of neural dynamics are *undesirable*, rather than discerning which features of the dynamical system are lacking. For example, tremors in Parkinson’s disease (PD) are associated with a characteristic beta oscillation (*i.e.*, 13-30 Hz) of the local field potential in the subthalamic nucleus, and state-of-the art feedback control strategies use this signal to trigger deep brain stimulation (DBS) until the beta oscillation subsides [Malekmohammadi et al., 2016]. Similar neural signatures are also present for epilepsy [Handforth et al., 1998, Morrell, 2011]. A model “diseased” system with three stable fixed points representing three possible voluntary movement command was constructed with an additional, unwanted spiral attractor representing the beta oscillation macrostate. Difficulty in initiating voluntary motion (bradykinesia) in PD patients could be due to strong attractive macrostate [Nini et al., 1995, Little and Brown, 2014]. Using myopic control, we manipulated the dynamics to match the target dynamics of a healthy system structured using the design principles from section 2.3 to avoid the avoid beta oscillation state while preserving the fixed points of the system. The phase portrait of the target dynamics are shown in Figure 6.

**Figure 6:**
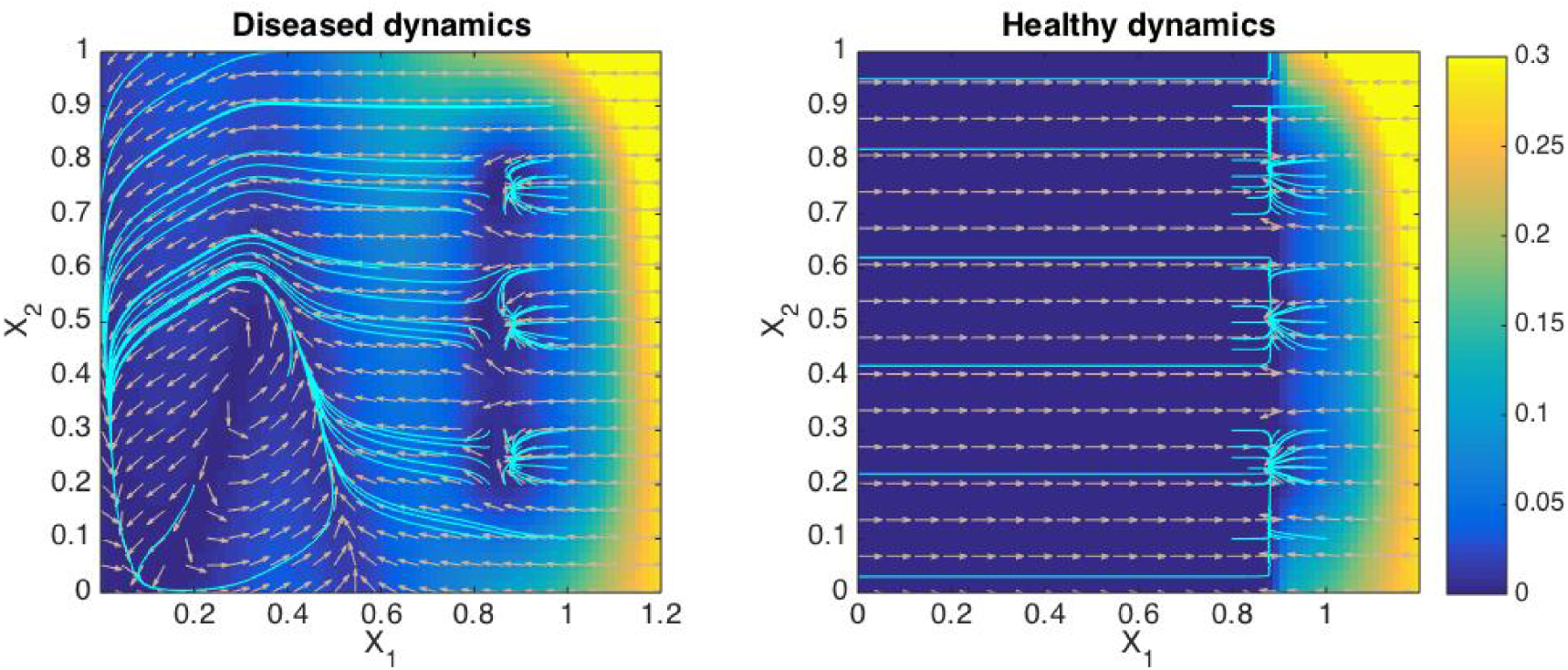
Phase portraits for diseased and healthy dynamical systems. Color denotes the magnitude of the dynamics ℱ and 𝒢, and the direction is shown by the arrows. Streamlines are shown in cyan. The diseased system contains three stable points, but also a spiral attractor at small values of *X*_1_ and *X*_2_. The target healthy dynamics has been designed to contain slightly repulsive dynamics in the original spiral attractor region, but still maintains its stable points.

The overall performance of myopic control is summarized in Fig. 7. A sample of 500 trials points was initialized in the asymptotic distribution of the PD limit-cycle attractor, and state estimation was performed for 100ms in the absence of control before the control was switched on. Monitoring 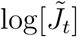 in Figure 7A, individual trials reflect the initial oscillatory behavior of being in the disease state before sharply declining, whereas the trial-averaged behavior shows the overall improvement due to control. This remarkable removal of disease-state behavior is further demonstrated in state-space trajectory of a typical trial (Fig. 7B). Once control is switched on the target dynamics successfully lead it out of the limit cycle and into a stable attractor point. A spectrogram of the state *X*_1_ for an analogous, longer simulation is shown in Fig. 7 D. There, a beta oscillation endured for 1s, and then myopic control was switched on to evolve to a stable point. The spectrogram reflects the oscillations during the uncontrolled period, and once the control is switched on it subsides and leaves only low-frequency components as it moves toward the stable point. The optimal control signal for the colored trajectory in Fig7A in shown in Fig. 7C. it is modest in amplitude relative to the magnitude of the dynamics, and has a straightforward waveform, demonstrating that given only minimal additional consideration to constraints on the control signal that myopic control could feasibly, efficiently, and safely be implemented in living subjects.

**Figure 7:**
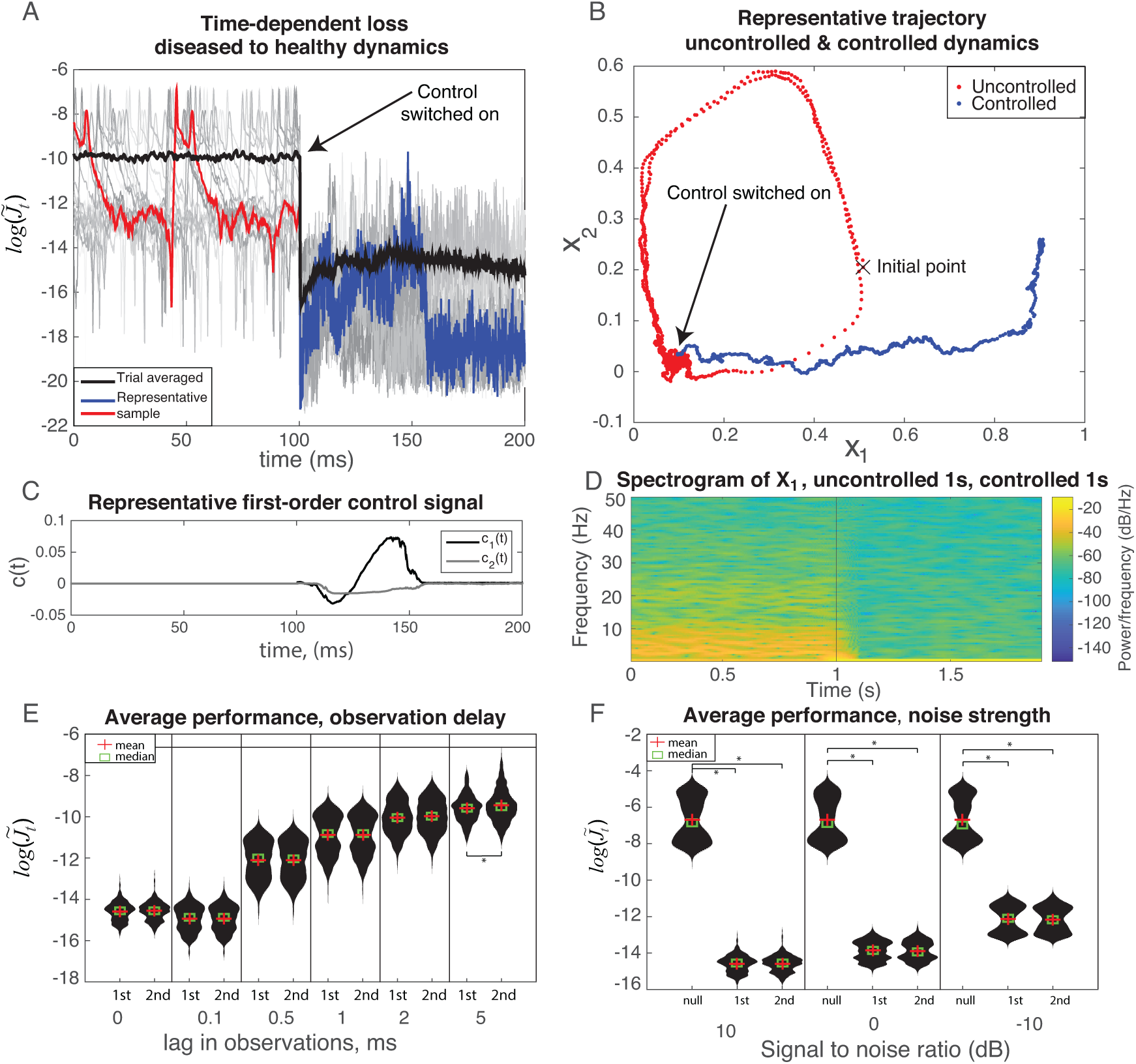
A) Time-dependent performance of first-order control for small observation noise (SNR 10dB). Instances of simulations are shown in gray, while the mean behavior is given in black. A representative trial has been singled out in blue and red for additional analysis. Initial trajectories are uncontrolled (blue), and allowed to fall into the asymptotic distribution of the limit cycle before control is switched on (red). Costs are plotted as time-locked 100ms before the control is switched on. B) Trajectory of typical trial through state space shown before and after the control is implemented, demonstrating a move back towards healthy state space. C) Control signal of a representative trial. D) Spectrogram of *X*_1_ for an analogous, longer simulation of uncontrolled evolution for 1s (noted by vertical line), and myopic control for final 1s. E) Violin plots showing distribution of time-averaged log-performance for a lag in observations, requiring state prediction. Horizontal line corresponds to the average log-loss of null control. F) Similar violin plots as in E), but for for varying observation noise strength, and including null control (*u*_*t*_ = [0, 0]). In both E) and F) * indicates *p <* 0.001 for a two-sample t-test. Unless otherwise noted, samples are not significantly different.

Finally, we benchmarked the performance of first- and second-order control as compared to uncontrolled dynamics by calculating distributions of the time-averaged log-cost 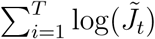 for varying lags between state observation and control signal calculation (Fig. 7E), and for different observation noise strength (Fig. 7F). While there is a interesting trend in the stretching of bimodal distribution into a near unimodal one at high observation lag, we observed no impactful difference between first- and second-order control with an increasing delay of observations. Similarly, there is a transition to a distinct bimodal distribution at large signal to noise, though both controllers perform similarly.

## 4 Discussion

Here we developed a perspective on what features are necessary for a flexible control of any dynamical system underlying neural computation. The controller should function to assist the dynamical system performing the computation, not taking on the role of a dynamical system, itself. In order for the controlled dynamics to function as a separate dynamical system on its own, we proposed a myopic control scheme that alternatively manipulates the dynamics to function as a set of target dynamics over a single time step, as opposed to trajectory-tracking controllers that function over a finite time horizon and must first perform the neural computation on their own. We developed an approximation of this control for nonlinear dynamics that is separable from state estimation, provided direction about design principles for how to construct a targeted dynamical system, and demonstrated its application in two varied scenarios. In both examples, first order control performed comparably to second-order control, showing the potential to generate feasible control signals that function under practical conditions.

The base of our controller formulation is reminiscent of the model-based control and model reference (adaptive) control (MRAC). Utilizing model-based control alongside quality state estimation [Watter et al., 2015] to manipulate neural dynamics is an attractive strategy that can harness machine learning methods to build effective, patient-specific statistical models of the brain by using real-time patient data, which could then be used as precision medical treatment [Collins and Varmus, 2015, Ozomaro et al., 2013]. The initial efforts of MRAC focused heavily on adaptive update rules for estimating the parameters of different forms of target plants (dynamics, in our work), and predominantly one adaptive controller form was utilized: strictly positive real (SPR) Lyapunov design. This form of controller depended on a SPR transfer function formulation of its plant dynamics, as was designed to guarantee bounded control signals that can track target trajectories or regulate to a fixed point from a target plant. Our controller structure is similar in form to SPR control and could benefit from the similar extensions that took place in MRAC, such as analysis of the Lyapunov stability to rigorously establish safe bounds on the control [Ioannou and Sun, 2012] and the use of neural networks capable of handling nonlinear plant dynamics [Narendra and Parthasarathy, 1990, Hagan et al., 2002].

This is not the first neural controller to consider neural variability as an important component to preserve in neural systems. Todorov and Jordan suggested a “minimal intervention principle” for neural systems that allows for deviations from a target trajectory, provided that they do not interfere with the target task [Todorov and Jordan, 2013]. The target was considered as a single point in state space, and their formulation allowed for high redundancy in the number of optimal trajectories that reached the target with the same cost. Their controller only corrects the trajectory when failing to act would result in a worse-than-optimal cost. While this is the only instance of control that acknowledges and respects neural variability during control, even prescribing a single point in state space as a target falls short of the general goals accomplished by myopic control to generate an entire target dynamics. For example, returning to the qualitative operation of myopic control in Fig. 1B, minimal intervention control would perform comparably to trajectory control by forcing state evolution in a non-dynamical fashion, while also restricting the neural variability that lead to an alternative fixed point.

An important feature of myopic control was its modular design. We studied a form in which only low-order moments of the state distribution were used in the controller, which decoupled the controller form from state estimation and allowed for any state estimator to be implemented. First-order control is considerably more straightforward to use because of its lack of higher-order derivatives on the dynamics, which may also come with a benefit being a more robust controller during practical instances in which the dynamics must be inferred from data. Operating in the perturbative regime 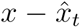 through regular state estimation small would ensure that first-order methods are successful.

A key issue that dictates myopic control’s qualitative success is the choice or design of target dynamics. Target dynamics could be designed by modifying the current dynamics through either addition, removal, or modification of specific features in the state space. There is an appeal to omitting features, as this approach resembles lesioning studies that aim to infer causal importance to behavior.

In our beta oscillation example, omitting the limit cycle appeared to be the safest and most practical approach. However, if more experimentally accurate understanding of the Parkinsonian dynamics suggests that omitting a limit cycle could introduce unwanted behavior, it may be more prudent to modify the limit cycle with an exit pathway. Still, one may wish to study the extent of a neural system’s computational flexibility by adding features as we did with our decision-making example.

It should be noted the adding in new features can be a tedious process in practice, as it took considerable parameter tuning to create our robust neural integrator system. Additionally, we considered only two-dimensional systems, but interesting dynamical systems may in fact lie in higher dimensions [Park et al., 2014]. Our design approach of filtering functions is general enough to extend to high dimensions, though implementing them in practice may take additional care. Removing features with smoothed versions of step functions would still work, as would adding stable points with gabor functions, though they would need to be high-dimensional analogues. Visualization and analysis tools for high-dimensional spaces could help determine the hypervolumes to omit, and how (an)isotropic the features must be.

Myopic control is certainly not the only form of controlled stimulation of neural systems, and it is important to note how these methods differ from our perspective. One of most successful applications of neural stimulation is in the field of neuroprosthetics, where implants mimic afferent sensory inputs such as cochlear or retinal implants, or translate efferent outputs into motor actions for artificial limbs [Sanchez, 2015]. The control strategies behind these technologies are complex and varied compared to myopic control [Schiff, 2011], though this is required in part because the goal of neuroprosthetics is distinctly different: the controlled (de)coding of these neural signals do not constitute a dynamical system, but rather interacts with the pre-existing normal neural dynamics of the area as inputs. Neural prosthetics for cognitive function, for example, memory processing in hippocampus [Berger et al., 2011], are much more amenable to myopic control scheme, since the normal function of the neural system constitutes a dynamical system.

Deep brain stimulation (DBS) for neurological disorders (*e.g.,* Parkinson’s disease) is a control application within the scope of myopic control, as we demonstrated with our first example study. A recent approach to DBS that harnesses neural recordings uses a model-free method to simply reduce beta-band oscillations seen in local field potential recordings in the basal ganglia [Malekmohammadi et al., 2016], a potential neural signal related to PD symptoms [Quinn et al., 2015, Trager et al., 2016]. The disadvantage to such a heuristic approach is that the link between beta oscillations in basal ganglia and cortex, let alone its relationship to actual PD symptoms, is still not fully understood. Moreover, other feedback targets are being actively considered as well [Rosin et al., 2011, Little and Brown, 2012]. Myopic control allows us to causally investigate the role of neural signatures correlated with the disease—we can specifically target fixes to the abnormal dynamics for beta oscillations, for example, and improve our understanding of the disease and also improve treatments.

Our first example was motivated from a position of understanding neural dynamical systems for evidence accumulation and decision making, and more generally to demonstrate its application as a tool to causally investigate cognitive processes. Several models of evidence accumulation have been considered in the context of using variability in spiking dynamics of lateral intraparietal cortex (LIP) in monkeys [Churchland et al., 2011, Huk and Shadlen, 2005, Resulaj et al., 2009], and one future experiment could attempt myopic control using different models for the control systems to produces a given target system, say RNI. Performing myopic control in that context would be a more powerful approach than perturbative, random stimulation of the system to simply infer parameters of an underlying dynamical model represented in LIP. Additionally, a more sophisticated experiment could attempt to utilize controlled stimulation to force the opposite decision of a target dynamics; the success of which would not only provide evidence that the controller is operating based upon the correct dynamical systems model, but would also constitute a substantial advance in the control of cognitive dynamics.

The history of advances in model reference adaptive control (MRAC) provides a strong template for how myopic controllers for neural dynamics control could be developed. Our work here assumed a known model for the controlled dynamics, and future work should integrate adaptive estimation of the controlled dynamics, themselves, into the controller. In particular, as an extension of the initial neural network structures used to perform MRAC [Narendra and Parthasarathy, 1990, Hagan et al., 2002], there is opportunity to utilize deep networks that accomplish adaptive estimation these dynamics and their states within a neural-network myopic controller architecture [Zhao and Park, 2018, 2017, Sussillo et al., 2017].

A larger and more immediate question is what steps must be taken to implement myopic control experimentally? The most important underlying component is access to quality neural measurements. That is why recent work combining neural stimulation and observation as in [Tan et al., 2018] is so vital. In our work we assumed that the ground-truth neural dynamics for both the controlled dynamics ℱ and the target dynamics 𝒢 were known, but in practice they must estimated from neural measurements. We demonstrated that first-order myopic control can function well, which necessitates estimation of only the state mean over higher order moments, but first order control also requires estimating the full dynamics in (18).

The timescale of the underlying neural computation also suggests practical consideration. Since myopic control is designed as an online control, the state estimations and estimation of the dynamics must be fast in order to implement in real time. Longer time constants for processes that are characterized by a smaller total dynamics 𝒢*_t_* lead to slower changes in neural state, which allow for more accurate online state estimation, and thus a more accurate control signal. Akin to a slower moving target in state space, the less the dynamics have progressed, the more up-to-date that state information will be, and the better the control performance for slower dynamical processes. This motivated our demonstration that myopic control can still function well with a lag between neural observations and control implementation.

Moreover, estimating latent state dynamics is a difficult task altogether [Breakspear, 2017], and would likely need to be performed prior to control use, with adaptive updates to the dynamics estimation occurring online. Taking into consideration a generic framework i) signal processing (*e.g.*, spike sorting), ii) control signal calculation, and iii) delivery of stimulation; it seems reasonable to assume ∼5 ms of time required for myopic control, which is comparable to other closed-loop control estimates [Ciliberti et al., 2018]. In this regime of time lags of less than 5 ms, myopic control was demonstrated to perform well, which is promising for its implementation.

## Acknowledgments

This work was supported by the Thomas Hartman Center for Parkinson’s Research (64249).

